# Bayesian model-averaging of parametric coalescent models for phylodynamic inference

**DOI:** 10.1101/2025.02.22.639697

**Authors:** Yuan Xu, Kylie Chen, Dong Xie, Alexei J. Drummond

## Abstract

Bayesian phylodynamic models have become essential for reconstructing population history from genetic data, yet their accuracy depends crucially on choosing appropriate demographic models. To address uncertainty in model choice, we introduce a Bayesian Model Averaging (BMA) framework that integrates multiple parametric coalescent models– including constant, exponential, logistic, and Gompertz growth–along with their “expansion” variants that account for non-zero ancestral populations. Implemented in a Bayesian setting with Metropolis-coupled MCMC, this approach allows the sampler to switch among candidate growth functions, thereby capturing demographic histories without having to pre-specify a single model. Simulation studies verify that the logistic and Gompertz models may require specialised sampling strategies such as adaptive multivariate proposals to achieve robust mixing. We demonstrate the performance of these models on datasets simulated under different substitution models, and show that joint inference of genealogy and population parameters is well-calibrated when properly incorporating correlated-move operators and BMA. We then apply this method to two real-world datasets. Analysis of Egyptian Hepatitis C virus (HCV) sequences indicates that models with a founder population followed by a rapid expansion are well supported, with a slight preference for Gompertz-like expansions. Our analysis of a single metastatic colorectal cancer (CRC) single-cell dataset suggests that exponential-like growth is plausible even for an advanced stage cancer patient. We believe this highlights that tumour subclones may retain substantial proliferative capacity into the later stages of the disease. Overall, our unified BMA framework reduces the need for restrictive model selection procedures and can also provide deeper biological insights into epidemic spread and tumour evolution. By systematically integrating multiple growth hypotheses within a standard Bayesian setting, this approach naturally avoids overfitting and offers a powerful tool for inferring population histories across diverse biological domains.

## Introduction

Understanding how biological systems change over time is central to many questions in biology, from population-level dynamics [1] to macroevolutionary patterns [2]. In evolutionary biology, such insights help clarify the processes underlying speciation events [3, 4], while in epidemiology, they illuminate the dynamics of rapidly evolving pathogens during viral outbreaks [5,6]. Population-level analyses are also increasingly important in tumour biology and somatic evolution, where cancer cell populations undergo mutation and selection [7–10]. The advent of single-cell sequencing [11] has transformed our ability to track these mutational events with unprecedented resolution, enabling detailed reconstructions of tumour evolutionary histories [12]. Together, these advances offer a flexible foundation for dissecting the origins, expansions, and adaptive trajectories of populations across diverse biological systems.

Phylodynamic models developed within a Bayesian framework provide a robust means to infer key dynamic parameters—such as growth rates and population sizes—from polymorphic gene sequence data. Approaches range from birth-death branching processes [13–16] to Kingman’s coalescent theory [17–19], each enabling rigorous reconstruction of historical population dynamics. Among these, coalescent-based models have emerged as standard tools in population genetics [20–22], simulating how changing population sizes shape genealogical relationships in randomly sampled lineages [17].

Classical coalescent theory initially assumed a constant population size, but subsequent extensions by Griffiths and Tavaré [23] and Donnelly and Tavaré [24] allowed for variable population sizes through integrable time-dependent functions. Rodrigo and Felsenstein [25] further broadened these applications to data sampled at multiple time points—such as rapidly evolving pathogens or ancient DNA—thereby expanding the scope of coalescent inference.

Modern coalescent inference can incorporate either non-parametric population functions, which flexibly adjust population size over multiple time intervals [26, 27], or parametric models that use specified mathematical functions to describe population trajectories [5, 28, 29].

Accurately estimating population size is critical for population genetic inference, yet it hinges on incorporating appropriate prior information within a Bayesian framework [30]. Ignoring such prior knowledge or misspecifying the population model can produce biased and unreliable inferences [31].

To advance Bayesian coalescent inference for single-cell sequencing data, we developed a Bayesian model averaging (BMA) approach that systematically addresses model uncertainty by integrating over a set of candidate models [32]. Unlike traditional model selection, which seeks a single best model, BMA acknowledges model uncertainty by combining multiple models’ predictions, weighting them by their posterior [33]. However, these model weights can be highly sensitive to prior specifications [34], particularly when comparing models of different complexity. We address this challenge through careful calibration of priors across our demographic models to ensure robust model averaging.

Biological populations frequently originate from a single founder individual, rapidly expanding until constrained by limited resources. This pattern characterises diverse scenarios, from viral spread within a host to the colonisation of new habitats by animals, and the proliferation of cancer cells [35–37]. While exponential growth aptly describes early population increases, it becomes untenable as resources diminish, ultimately giving way to slowed expansion and eventual stabilisation [38, 39]. The Gompertz model captures this transition particularly well [40]. Proposed by Benjamin Gompertz in 1825 as a tool to relate mortality and age [41], it quickly gained traction in insurance mathematics and has since been applied to a wide range of biological contexts, including mammalian [42], microbial [43], and avian growth [44, 45]. In tumour growth modelling, the Gompertz model’s nonlinear, sigmoidal curve mirrors the characteristic stages of neoplastic development: initial slow growth, subsequent rapid expansion, and eventual deceleration as nutrient availability, immune responses, and spatial constraints impose biological and physical limits on further proliferation.

In this study, we extend the coalescent inference framework by incorporating a suite of parametric population growth models—constant, exponential, logistic, and Gompertz—alongside their “expansion” variants, into the BEAST2 [46] phylogenetic software platform. After verifying the calibration and performance of these integrated models using simulated data, we leveraged a composite model space to implement Bayesian model averaging. This BMA approach systematically addresses model uncertainty, enabling robust inference of historical population dynamics under a wide range of demographic scenarios. By further validating the BMA framework on simulated datasets and subsequently applying it to empirical data, we demonstrate that our enriched modelling strategy not only enhances the accuracy and stability of parameter estimates but also provides deeper insights into the evolutionary processes governing complex biological populations.

The remainder of this paper is organised as follows. First, we describe our implementation of growth models and BMA framework in BEAST2. We then validate our approach through simulation studies, demonstrating proper calibration and mixing. Finally, we apply our method to HCV and CRC datasets, revealing distinct population dynamics in these biological systems.

## Methods

### Inference framework

The coalescent model provides a flexible framework for relating observed genetic variation in a sample to underlying population demographic history [22]. By describing the genealogical relationships among sampled sequences under a probabilistic model, it can accommodate any deterministic, time-varying population size function *N* (*t*) [23]. Accurate estimation of *N* (*t*) is particularly important in epidemiological and population-genetic studies, where incorporating detailed prior information via a Bayesian framework can substantially improve inference accuracy.

Our phylodynamic inference is grounded in the joint posterior distribution:

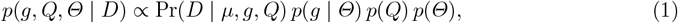

where *D* denotes the observed sequence alignment, *µ* is the molecular clock rate, *g* represents the phylogenetic time tree, *Q* comprises substitution model parameters, and *Θ* encodes the parameters of the population-size function *N* (*t*). The alignment *D* is assumed to evolve according to a continuous-time Markov process along the branches of *g* [47], and both the tree and population dynamic parameters are jointly estimated from the sequence data.

Consider a rooted binary time tree *g* estimated from *n* contemporaneously sampled sequences, with the sampling time defined as *t* = 0. Let *T* = {*t*_1_, *t*_2_, …, *t*_n−1_} be the set of coalescent times (moving backward in time) and *Δt*_i_ = *t*_i_ − *t*_i−1_ the intervals between successive coalescent events. In this setup, each coalescent event reduces the number of extant lineages by one. While we focus on contemporaneous sampling here, extensions to serial sampling are well-developed [25, 30, 48].

For a given number of lineages *k*_i_ at time *t*_i_, the instantaneous coalescent rate is *f* (*t*_i_) = 1*/N* (*t*_i_), where *N* (*t*_i_) is the population size at that time. Integrating this rate over time yields the cumulative coalescent intensity:

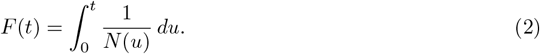

Because the coalescent waiting times are exponentially distributed, the probability density for observing a coalescent event at time *t*_i_, given that the previous event occurred at time *t*_i−1_, is:

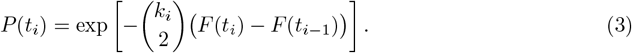

By combining these terms over all intervals, the probability density of the genealogy *g* given *Θ* can be expressed as:

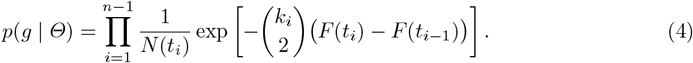

Provided that 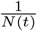 is integrable over time, this coalescent density can be computed for any specified demographic function *N* (*t*).

In practice, simulations often rely on inverse transform sampling to generate coalescent times from a specified *N* (*t*). Defining the inverse of the cumulative intensity function *F* ^−1^(*x*) as the time *t* at which *F* (*t*) = *x*, uniformly distributed samples in *x*-space can be mapped to coalescent times in *t*-space, thus reconstructing historical population size trajectories from genetic data. When analytical solutions for *F* ^−1^(*x*) are unavailable, numerical methods provide a viable alternative.

### Deterministic population functions

This study applies four mathematical growth models to phylodynamic inference: the constant population model, the exponential growth model, the logistic growth model, and the Gompertzian growth model. The latter three functions can be described in two forms: a base function, representing population dynamics in the absence of an ancestral population size (*N*_A_ = 0), and an expansion function, obtained by introducing a non-zero *N*_A_. This expansion accounts for ancestral contributions and is better suited for scenarios involving founding populations or prior population bottlenecks.

The exponential model provides a clear example. Its baseline form is given by:

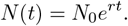

By introducing a non-zero ancestral population size *N*_A_, we obtain the exponential expansion function:

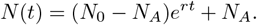

When *N*_A_ = 0, this expansion form reduces to the baseline model, indicating that the key difference lies in the incorporation of the ancestral population component.

The logistic model, characterised by parameters *K, b*, and *t*_50_, offers a sigmoidal representation of population growth. Its baseline form is:

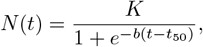

where *K* is the carrying capacity, *b* governs the rate of growth, and *t*_50_ specifies the time at which the population reaches half of *K*.

Incorporating an ancestral population size *N*_A_ adjusts the model to:

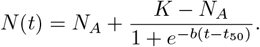

This variant preserves the logistic shape but shifts the baseline to *N*_A_. The parameter *t*_50_ still denotes the time at which the population is halfway between *N*_A_ and *K*. Introducing *N*_A_ thus extends the logistic model to scenarios involving non-zero ancestral populations.

The Gompertz model, characterised by an exponentially decreasing growth rate, is widely used to describe tumour growth dynamics. Among the various parameterisations proposed in the literature, the Gompertz-Laird formulation [49] is frequently adopted. This classical representation involves three parameters: the initial population size *N*_0_, the growth rate parameter *b*, and the carrying capacity *N*_∞_. Its baseline (no ancestral population) form is

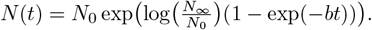

To account for a non-zero ancestral population, we introduce a Gompertz expansion:

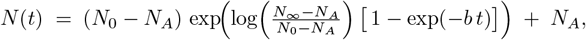

which reduces to the baseline form if *N*_A_ = 0. Here, *N*_A_ shifts the growth trajectory upward so that the population grows from an existing size rather than near zero.

Although the parameters {*N*_0_, *N*_∞_, *b*} are biologically intuitive, *N*_0_ and *N*_∞_ can be strongly correlated in MCMC sampling, potentially slowing convergence. To address this issue, adaptive methods such as the Adaptable Variance Multivariate Normal (AVMN) operator [50] can be employed to learn and adjust parameter correlations during the MCMC run.

Additionally, alternative parameterisations can be considered to further reduce correlation and enhance computational performance. One approach is to define 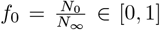. This ratio *f*_0_ is dimensionless and constrained to the interval [0, 1], enabling the straightforward use of a Beta prior without directly depending on the absolute sizes *N*_0_ or *N*_∞_. Another approach, mirroring the logistic model, is to replace *N*_0_ with a half-capacity time *t*_50_ and *N*_∞_ such that 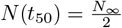 in the baseline case, or 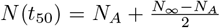 when an ancestral size *N*_A_ is present. This {*N*_∞_, *b, t*_50_} parameterisation helps align Gompertz with logistic growth, making *t*_50_ a consistent reference time in both models. Detailed derivations of these transformations and prior considerations can be found in *Supplementary 1*.

Although in this part we initially introduced the logistic model via {*K, b, t*_50_} and the Gompertz model via {*N*_∞_, *b, t*_50_}, in practice these parameters often serve similar roles (for example, *K* and *N*_∞_ both describe an asymptotic population size). In some contexts, one may further rename them into a single “maximum size” parameter (e.g., *N*_max_) if it is more convenient to treat them as conceptually equivalent.

### Model selection and averaging

As described in the previous method section, our Bayesian framework infers the joint posterior over the genealogy *g*, the substitution model parameters *Q*, and the demographic parameters *Θ*. We now extend *Θ* to include the discrete indicators *I* and *I*_A_. Specifically, we define

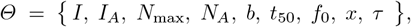

where *I* selects one of the growth-function families, and *I*_A_ ∈ {0, 1} determines whether a non-zero ancestral population size *N*_A_ is included. *x* and *τ* are the two times separating the phases of the C–E–C piecewise population function (See Section 1 for further details on these piecewise formulations.) The prior on *Θ* includes uniform distributions for *I* and *I*_A_, letting the data inform which demographic model is favoured and whether an ancestral component is warranted.

To unify notation, we introduce *N*_max_ as the principal population-size parameter across all models. In the constant and exponential formulations, *N*_max_ represents the initial (or constant) population size, whereas in the logistic and Gompertz models, it serves as the carrying capacity (i.e., the upper bound). Although Gompertz growth can alternatively be parameterised via {*N*_0_, *f*_0_, *b*}, we do not employ both parameterisations in the same analysis; thus, for consistency, we use a single symbol, *N*_max_, to denote the population-size parameter for all models.

Depending on the biological context, one may either unify the logistic parameter *K* and the Gompertz parameter *N*_∞_ into a single *N*_max_ if they are considered the same conceptual quantity, or retain them as separate parameters if their mechanistic interpretations differ. Likewise, one may choose to share a same prior for *N*_max_ across multiple models, or instead assign distinct priors to reflect uncertainty or variation in how each model handles the asymptotic population size. Similarly, growth-rate parameters can be treated as separate or identically distributed. When parameters in different models have the same prior, then sharing that parameter is conceptually identical to having distinct parameters, but can improve overall mixing by reducing the dimensionality of the combined model space.

To provide a unified view of how these parameters are employed across the various model variants, we summarise in Table 1 the usage of each parameter for all (*I, I*_A_) combinations. In the table, a tick indicates that the parameter is *used* in that model variant, whereas a empty cell indicates that the parameter is *not used*. Green rows indicate model variants that are included in the model averaging for that data set. Gray rows are models that are not considered. Note that constant-size models (*I* = 0) inherently have no expansion form (thus *I*_A_ is irrelevant), whereas exponential, logistic, and Gompertz functions each appear in baseline (*I*_A_ = 0) or expansion (*I*_A_ = 1) form. For Gompertz, we use either an *f*_0_-based or *t*_50_-based parameterisation, but not both simultaneously. Finally, the parameters *x* and *τ* appear only in the piecewise models that involve a change point.

**Table 1:**
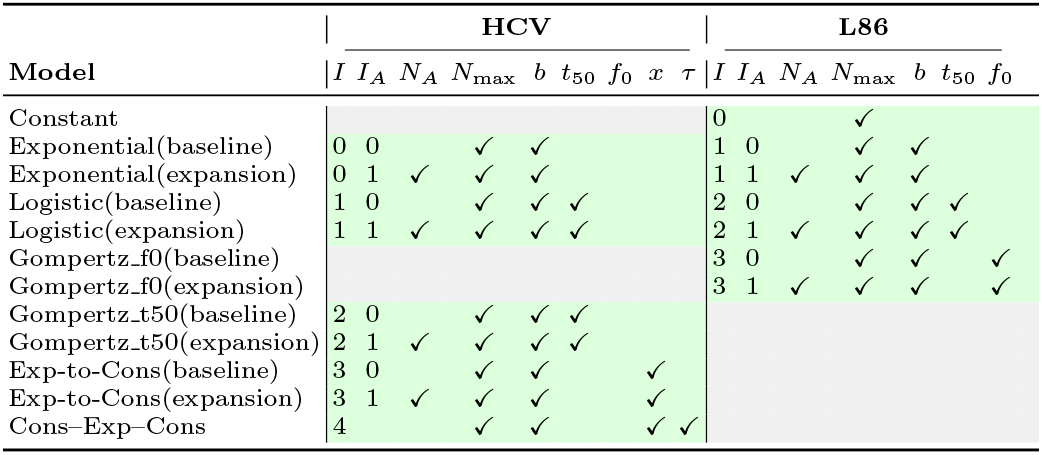
Parameter usage across different model configurations. This table illustrates which parameters in {*I, I*_A_, *N*_max_, *N*_A_, *b, t*_50_, *f*_0_, *x, τ*} are active (green) or inactive (gray) under each combination of the discrete indicators (*I, I*_A_). For illustration, we list the models used in the HCV and L86 analyses, respectively.

We will employ these models in both the HCV and L86 applications (see below), but present this consolidated table here so that the reader can see the complete set of parameter configurations at a glance before we move to real data analyses. For details of the implementation, please refer to the scripts in Supplementary Code S.1 and S.2.

Although these indicators can be updated via standard MCMC steps, high-dimensional or multimodal posterior distributions can challenge efficient mixing and increase the risk of becoming trapped near local maxima. To mitigate these issues, we additionally employ Metropolis-coupled MCMC (MC^3^) [51], which runs multiple chains in parallel at different temperatures, together with an adaptive strategy for tuning temperature differences [52]. This approach helps maintain effective acceptance rates and improves overall mixing performance.

To avoid bias, both indicators, *I* and *I*_A_, are assigned uniform priors, thus letting the data determine which demographic model is favoured and whether the ancestral phase improves model fit. This approach not only enables Bayesian model selection by identifying which (*I, I*_A_) combination attains the highest posterior support, but also facilitates model averaging, in which inferences about population trajectories or tree topology are weighted by each model combination’s posterior probability.

### Model validation and simulations

To assess the reliability and calibration of our Bayesian coalescent models and the model-averaging approach, we conducted a series of simulation-based evaluations. Previous studies indicate that coalescent models assuming constant or exponentially changing population sizes are generally well-calibrated. Building on this foundation, we focus here on the logistic and Gompertz growth models (which each have multiple parameterisations) and evaluate whether our Bayesian model averaging (BMA) framework maintains robust inference under these more complex demographic scenarios.

We used the LPhyStudio and LPhyBEAST packages within the LinguaPhylo framework [53] to simulate data under each demographic model, generating 100 replicate datasets per model. To vary the amount of phylogenetic signal, we considered two sample sizes (*n* = 16 and *n* = 40) and two sequence lengths (200 bp vs. 2000 bp). This creates settings with lower or higher evolutionary information, providing a more stringent test of parameter estimation and model recovery.

We explored two substitution models, JC69 and GT16 [54, 55]. JC69, with uniform rates and nucleotide frequencies, served as a baseline scenario, whereas GT16 incorporated a Dirichlet prior on nucleotide frequencies and substitution rates, emulating greater complexity often seen in somatic evolutionary processes.

All simulated datasets were analysed in BEAST2 using MCMC with sufficiently long run times to ensure convergence (monitored via ESS values and trace plots). When strong correlations among parameters were detected (e.g., in Gompertz models), we introduced additional sampling operators (e.g., up-down operators) to facilitate mixing and faster convergence. Table 2 summarises the key parameters and their prior distributions for the logistic and Gompertz models.

**Table 2:** Prior distributions for parameters under each demographic model. For all LogNormal distributions, *µ* (meanlog) and *σ* (sdlog) refer to mean and standard deviation on the log-scale. For the Beta distribution, *α* and *β* are shape parameters.

|  | Logistic | Gompertz (t50) | Gompertz (f0) |
| --- | --- | --- | --- |
| **Parameter 1** | $t_{50}$ | $t_{50}$ | $N_0$ |
| *Distribution* | LogNormal($\mu = 0.3$, $\sigma = 0.1$) | LogNormal($\mu = 0.3$, $\sigma = 0.1$) | LogNormal($\mu = 8$, $\sigma = 0.5$) |
| **Parameter 2** | $b$ | $b$ | $b$ |
| *Distribution* | LogNormal($\mu = 2$, $\sigma = 0.5$) | LogNormal($\mu = 2$, $\sigma = 0.5$) | LogNormal($\mu = -0.95$, $\sigma = 0.2$) |
| **Parameter 3** | $n\text{CarryingCapacity}$ | $N_\infty$ | $f_0$ |
| *Distribution* | LogNormal($\mu = 5$, $\sigma = 0.3$) | LogNormal($\mu = 4.5$, $\sigma = 0.5$) | Beta($\alpha = 20$, $\beta = 7$) |

### Data analysis

In addition to the simulation study, we applied our BMA framework to two real-world datasets: (1) Egyptian Hepatitis C virus (HCV) sequences, and (2) single-cell genomic data from the L86 metastatic colorectal cancer (CRC) study. While the same MCMC-based approach was used for both, each dataset involved a slightly different set of candidate models to suit its biological context. In the following subsections, we describe which demographic functions and expansions were considered for each case. We performed MCMC until convergence, then examined the posterior model probabilities to identify which demographic assumptions received the strongest support.

### Parametric posterior-based demographic reconstruction

After completing each MCMC run (from real data), we reconstruct historical population sizes from the model-averaged posterior on a set of discrete time points. Each iteration of the Markov chain yields a demographic model indicator and its associated parameters, defining a specific population-size function. Once sampling is finished, we define a suitable time grid—for example, from the present to the 95% HPD upper bound of the tree height—and, for each posterior sample, compute *N* (*t*) across these lattice points. We then compile these values into a posterior distribution of population sizes for each time point. Summaries such as mean, median, and credible intervals (e.g., 95% HPD) can be derived from these distributions. Plotting these statistics against time produces a continuous population trajectory that reflects both parameter and model uncertainty: if certain parametric forms appear more often in the chain, their trajectories naturally receive greater weight in the final estimate.

### Implementation

All methods described in this study have been implemented in an open-source software package named PopFunc, which is hosted in a public GitHub repository ^1^ This repository contains the core source code, modularized components for various population growth models (including Gompertz and logistic expansions), and a plugin system compatible with our Bayesian inference framework. A step-by-step example tutorial is provided in the same repository’s README, illustrating the entire analysis workflow—from an initial demo template, to generating an MC^3^-compatible BEAST2 XML, offering a practical guide for reproducing our methods.

To ensure transparency and reproducibility, we maintain a separate repository titled PopFunc-paper ^2^, where we provide all simulation scripts, configuration files, and real-data analysis pipelines corresponding to this manuscript. In particular, PopFunc-paper includes:

- Example scripts to replicate the simulation studies in Section 1, along with instructions for parameter settings.
- Data-processing workflows for the Egyptian HCV and L86 CRC datasets (Section 1), including raw input files and post-processing routines to generate all figures and tables.

Both repositories are released under the MIT License. The scripts and code therein are sufficient to reproduce all analyses and results reported in this paper.

## Result

### Evaluation on Simulated Datasets

We first assessed the performance of the Logistic coalescent model by monitoring MCMC convergence and parameter estimation in Tracer. Across 100 replicate simulations, the average effective sample size (ESS) values for all parameters exceeded 200 (see *Supplementary Figure S1* for detailed calibration plots).

We then applied the same chain-length settings to the Gompertz Coalescent model, testing two parameterisations. In contrast to the Logistic model, the Gompertz model exhibited lower ESS, suggesting strong parameter correlations that hinder exploration (see *Supplementary Figure S2*). Further joint-marginal analyses of a representative replicate revealed strong parameter correlations, which hindered efficient exploration when using standard independent proposal operators.

To address these challenges, we introduced up-down operators and integrated the Adaptive Variance Multivariate Normal (AVMN) operator [50], which learns the underlying posterior correlations and adapts the proposal distribution during the MCMC run. Implementing these strategies substantially enhanced sampling efficiency for the Gompertz model, with ESS values demonstrating a marked improvement across all parameters (see *Supplementary Figure S3*). In particular, after applying specialized operators, 91%–99% of the true values fell within the estimated 95% HPD intervals, indicating a well-calibrated method.

Next, we examined a more complex substitution and error model, the GT16 model, which accounts for amplification and allelic dropout errors in single-cell sequencing [54, 55]. Despite the increased complexity, the combination of correlated-move operators and the AVMN operator maintained good chain mixing and ESS values (see *Supplementary Figure S4*).

Additionally, we performed analogous simulations for the expansion variants (non-zero ancestral population) of the exponential, logistic, and Gompertz models, which also demonstrated well-calibrated parameter estimates (see *Supplementary Figures S5–S8*).

Finally, to assess how effectively our model-averaging approach can recover the correct population function under a broader set of demographic assumptions, we conducted a simulation study using 40-taxon trees (500 bp per sequence) under a strict molecular clock and the HKY substitution model. We considered four demographic families—Constant, Exponential, Logistic, and Gompertz—and included both baseline and expansion variants for each, except for the Constant model which, by definition, does not have an expansion form. This yielded a total of seven candidate models, all of which had previously passed our calibration tests (see the Supplementary for details). Here, we shared the same prior distributions on three key parameters, {*N*_A_, *N*_max_, *b, t*_50_}, across the relevant models. The BMA sampler yielded well-calibrated estimates for continuous parameters such as tree height, tree length, and the demographic hyperparameters. All simulation scripts, including the specified priors, are provided in the Supplementary code S.3.

Moreover, the BMA sampler successfully identified the true generating model with high coverage across 100 replicate analyses. Specifically, we summarizes the coverage metrics for the discrete model indicator *I*. Coverage for the indicator is the proportion of replicate analyses in which the true model is included in the 95% credible set. We find that the correct model is recovered in 98% of replicates (only 2 out of 100 did not include the true model in the 95% credible set). Moreover, the average posterior probability assigned to the true model is approximately 0.70, and the average of the highest posterior probability across all models is around 0.80, indicating that the correct model typically receives a substantial share of posterior support. Full simulation scripts, including the specified priors, are provided in the Supplementary code S.3.

### Hepatitis C Virus in Egypt

Hepatitis C virus (HCV) is genetically diverse and constitutes a leading cause of chronic liver disease worldwide. In particular, Egypt exhibits an exceptionally high prevalence of HCV, commonly attributed to large-scale parenteral antischistosomal therapy (PAT) campaigns during the mid-20th century [56–58]. At that time, unsafe injection procedures provided an efficient route for HCV transmission, facilitating rapid viral spread among the Egyptian population [59]. Pybus et al. [60] introduced a three-phase demographic model - constant–exponential–constant (C–E–C) - to characterize the transition from a stable HCV population size, through a phase of exponential growth concurrent with PAT, and finally returning to a high-level plateau. However, previous inferences often relied on rigid demographic assumptions imposed by fixed, piecewise population-size trajectories. Recent methodological advances now allow the simultaneous integration of phylogenetic, substitution, and demographic parameters within a Bayesian framework, enabling more flexible exploration of HCV’s historical expansion in Egypt.

We configure our Bayesian Model Averaging (BMA) framework with a set of parametric and piecewise demographic functions, excluding the purely constant-size model (which was deemed biologically implausible). Specifically, we incorporate:

- Baseline and expansion variants:
- We retain logistic, Gompertz, and exponential growth functions, each optionally combined with a non-zero ancestral population (*I*_A_ = 1), which introduces the parameter *N*_A_.
- Pybus’s C–E–C model: We include the original constant–exponential–constant framework from Pybus et al. [60].
- A new “Exponential-to-Constant” model:

This piecewise function transitions from exponential (or exponential expansion if *I*_A_ = 1) to a stable plateau at *N*_max_. In essence, it captures a trajectory similar to the C–E–C pattern but relies on fewer parameters. At time *τ*, the model switches from exponential growth to a constant size *N*_max_, thereby representing both the early rapid-rise phase and the subsequent plateau in a simplified manner.

All these models reside in a common parameter space, where *I*_A_ toggles the presence of *N*_A_ across exponential, logistic, and Gompertz functions, while the piecewise exponential-to-constant and C–E–C models each have their own “split time” plus *N*_max_. We impose a shared lognormal prior on *N*_A_ for all expansion variants, and unify logistic/Gompertz carrying capacities under a single lognormal prior for *N*_max_. The growth-rate parameter *b* in Gompertz has a separate prior from that of logistic/exponential, and both logistic and Gompertz share the same prior on *t*_50_. All these priors are made sufficiently broad, thus ensuring that the MCMC can freely explore the parameter space.

By running a Metropolis-coupled MCMC across this expanded model set, we can directly compare whether the data favor Pybus’s original C–E–C, a purely parametric function (e.g., Gompertz expansion), or the new exponential-to-constant design. For each posterior sample in the MCMC chain, we record which parametric model (*I, I*_A_) was selected, extract the relevant parameters (e.g., growth rate *b*, carrying capacity *N*_max_, ancestral size *N*_A_), and evaluate the population trajectory at a grid of time points. By aggregating these population-size values across all samples, we derive the posterior distributions at each time bin, from which we compute the mean, median, and 95% highest posterior density (HPD) intervals.

Table 3 presents the posterior probabilities and relative log Bayes factors (logBF) for each candidate demographic model under our Bayesian Model Averaging (BMA) approach. Notably, the *Gompertz Expansion* variant attains the highest posterior probability (0.438), emerging as the best-supported model. Two additional models - *Exponential expansion to constant* and *Logistic Expansion* - had moderate support (0.328 and 0.235, respectively). In contrast, the baseline versions of these functions (e.g., *Exponential, Logistic, Gompertz t*_50_), along with the piecewise *constant– exponential–constant* (C-E-C) model, had negligible support (i.e., their posteriors rounded to 0.00).

**Table 3:**
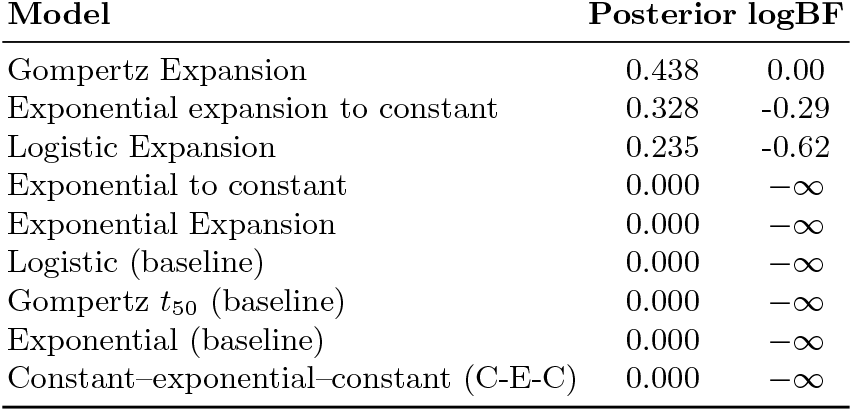
Posterior probabilities and log Bayes factors (logBF) for candidate models in the HCV dataset.

This outcome underscores the data’s strong preference for expansion-type models - those allowing for an ancestral population or additional growth-phase flexibility - over simpler or strictly piecewise assumptions. Models assigned a posterior of 0.00 also exhibit logBF values of −∞, indicating negligible support relative to the best model. Overall, these results highlight that incorporating an early founder population and subsequent rapid growth phase is critical for capturing Egypt’s HCV epidemic history.

Figure 1 illustrates the BMA-based demographic reconstruction of HCV in Egypt, revealing a pronounced expansion during the early 20^th^ century and a subsequent deceleration in later decades. The solid blue curve denotes the mean effective population size at each time bin, the dashed orange curve marks the median, and the shaded region encompasses the 95% highest posterior density (HPD) interval. These results are consistent with epidemiological data indicating that widespread intravenous treatment for schistosomiasis - implemented from the 1920s to the 1950s - likely triggered the marked rise in HCV infections [60].

**Fig. 1:**
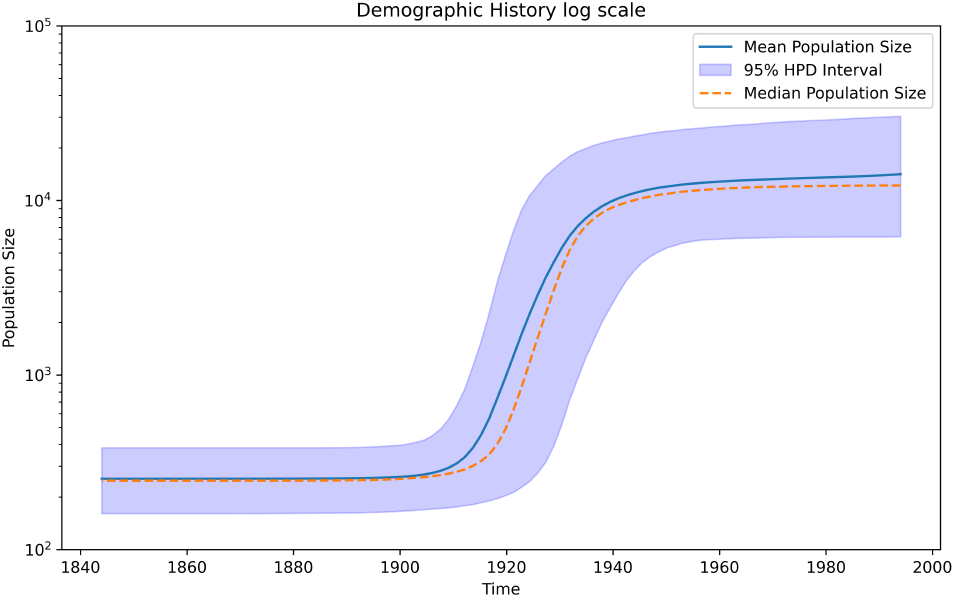
Parametric BMA Reconstruction of Egyptian HCV. The x-axis denotes calendar years, and the y-axis shows the effective population size on a log scale. The solid blue line is the posterior mean, the dashed orange line is the median, and the shaded region represents the 95% HPD interval. A pronounced rise in effective population size between the 1920s and mid-century is consistent with the era of large-scale parenteral antischistosomal therapy (PAT).

### Population Dynamics in the L86 Metastatic Colorectal Cancer Dataset

To further demonstrate our model-averaging method we investigate the metastatic evolution of colorectal cancer (CRC), analysing the L86 dataset. This dataset originates from single-cell DNA sequencing of a late-stage CRC patient who developed liver metastases [7]. Despite representing an advanced clinical scenario where tumour cells have already spread from the primary (colon) to the secondary site (liver), key questions remain regarding the subclonal architecture and local population dynamics at the metastatic lesion. Focusing on the metastatic subset can reveal the growth trajectories of subclones within the metastatic niche.

Following the GT16 substitution and error model [54], we configured a single-run Bayesian Model Averaging analysis to compare seven demographic models. Specifically, we included a simpler constant model and three major growth functions (exponential, logistic, and Gompertz), each allowed to appear in either baseline (*I*_A_ = 0) or expansion (*I*_A_ = 1) form. Note that for Gompertz, we specifically adopt the *f*_0_-based parameterization.

Table 4 summarizes the posterior probabilities and log Bayes factors (logBF) for the sampled models under this unified BMA scheme. Notably, the Exponential (baseline) model achieves the highest posterior probability (0.769), followed by Gompertz (f0) at 0.205. Together, these two account for over 97% of the total model support. In contrast, Gompertz (f0) Expansion, Logistic, and Exponential Expansion collectively account for under 3%. The Constant model had negligible posterior probability.

**Table 4:**
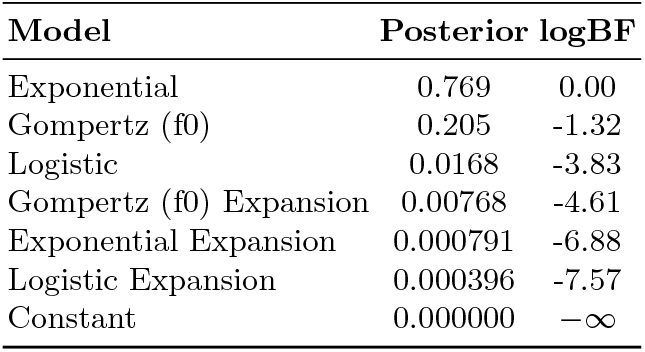
Posterior probabilities and log Bayes factors (logBF) for candidate models in the L86 metastatic CRC dataset under a single BMA analysis.

Figure 2 presents the BMA-based reconstruction of the metastatic population trajectory, pooling samples from all candidate models (both baseline and expansion). The solid blue curve indicates the posterior mean effective population size at each time bin, the dashed orange line denotes the median, and the shaded region encompasses the 95% highest posterior density (HPD) interval. A notable feature is the apparent capacity for continued or near-exponential growth, perhaps reinforcing the idea that metastatic tumours can retain proliferative potential even at advanced clinical stages.

**Fig. 2:**
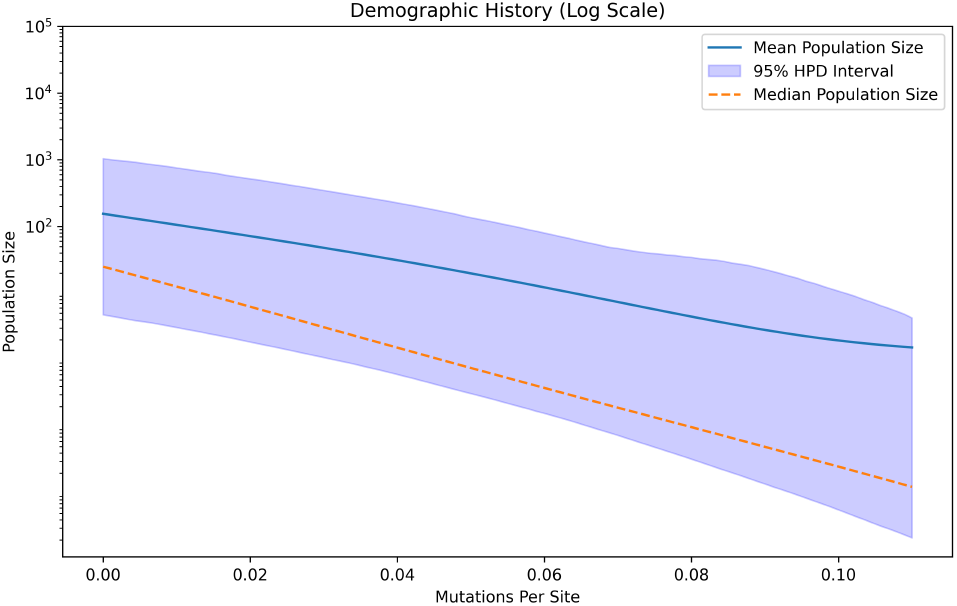
Parametric BMA Reconstruction of L86 Metastatic CRC. The x-axis represents relative time or approximate time since metastatic onset (scale is mutations per site), and the yaxis displays the effective population size on a log scale. The solid blue line is the posterior mean, the dashed orange line is the median, and the shaded region depicts the 95% HPD interval. Despite the advanced stage, the evidence suggests continued expansion in a near-exponential fashion.

These results suggest that tumour cells in the metastatic lesion may still be experiencing relatively rapid growth (as reflected by the Exponential and Gompertz (f0) baseline models), contrary to the intuition that a late-stage cancer would exhibit a plateau. The posterior probability of a non-zero ancestral population size was Pr(*I*_A_ | *D*) = 0.0089. Thus an ancestral population is not needed to explain this data.

## Discussion

In this work, we present an integrated Bayesian Model Averaging (BMA) framework that unifies multiple demographic models—including the Gompertz class—within a single coalescent inference scheme. By using discrete model indicators and Metropolis-coupled MCMC, our approach enables simultaneous exploration of alternative population-growth hypotheses, thereby mitigating the risk of overly committing to any single demographic function. This is a critical advantage in settings where the true growth dynamics remain uncertain or where existing models may impose overly simplistic or rigid assumptions.

A key methodological contribution of our study is the explicit integration of ancestral population expansions in the form of non-zero *N*_A_ terms. Whereas many classical coalescent analyses rely on baseline models such as constant or strictly exponential growth, we demonstrate that more flexible functions—e.g., Gompertz or logistic expansions—can be incorporated without sacrificing the Bayesian rigor of parameter estimation. Indeed, the BMA framework penalises overparameterisation automatically, only favouring additional complexity when supported by the data. This allows investigators to entertain a broad set of competing hypotheses, from purely exponential to logistic or Gompertz trajectories with ancestral population components, all within one unified inference run.

Here, our focus has been to demonstrate the feasibility and flexibility of BMA in phylodynamic inference, rather than to draw final conclusions about each empirical dataset. Consequently, our HCV and CRC examples serve primarily as methodological illustrations: they highlight how a unified BMA approach can recover plausible demographic histories under real-world conditions but do not attempt to establish a definitive epidemiological or clinical consensus.

In future work, researchers applying our framework to specific systems may wish to evaluate the impact of different priors more systematically, exploring how robust model weights and parameter estimates remain across a range of reasonable prior settings. Such sensitivity checks can help clarify whether the resulting inferences are driven by genuine signal in the data or by implicit assumptions about population growth. Moreover, although we have demonstrated the viability of our approach with two representative datasets, it could be further extended to incorporate additional demographic structures (e.g., multi-phase or bayesian skyline) or applied to larger singlecell genomic cohorts. In these more complex settings, the flexibility of Bayesian model averaging becomes especially advantageous, as it can reduce the risk of overfitting any single demographic function while still exploiting the strengths of each candidate model.

## Footnotes

1 https://github.com/LinguaPhylo/PopFunc

2 https://github.com/yxu927/PopFunc-paper

## References

1. Ronald Aylmer Fisher. The genetical theory of natural selection. The Clarendon press, Oxford, 1930.

2. Ernst Mayr. Systematics and the origin of species from the viewpoint of a zoologist, volume No. XIII. Columbia University Press, New York, 1942.

3. Tanja Stadler. Recovering speciation and extinction dynamics based on phylogenies. Journal of Evolutionary Biology, 26(6):1203–1219, 2013.

4. R Alexander Pyron and Frank T Burbrink. Phylogenetic estimates of speciation and extinction rates for testing ecological and evolutionary hypotheses. Trends in ecology & evolution, 28(12):729–736, 2013.

5. Oliver G Pybus, Michael A Charleston, Sunetra Gupta, Andrew Rambaut, Edward C Holmes, and Paul H Harvey. The epidemic behavior of the hepatitis C virus. Science, 292(5525):2323–2325, 2001.

6. Nuno R Faria, Andrew Rambaut, Marc A Suchard, Guy Baele, Trevor Bedford, Melissa J Ward, Andrew J Tatem, Joao D Sousa, Nimalan Arinaminpathy, Jacques Pépin, et al. The early spread and epidemic ignition of HIV-1 in human populations. Science, 346(6205):56–61, 2014.

7. Marco L Leung, Alexander Davis, Ruli Gao, Anna Casasent, Yong Wang, Emi Sei, Eduardo Vilar, Dipen Maru, Scott Kopetz, and Nicholas E Navin. Single-cell dna sequencing reveals a late-dissemination model in metastatic colorectal cancer. Genome Research, 27(8):1287–1299, 2017.

8. Nicholas Williams, Joe Lee, Emily Mitchell, Luiza Moore, E Joanna Baxter, James Hewinson, Kevin J Dawson, Andrew Menzies, Anna L Godfrey, Anthony R Green, et al. Life histories of myeloproliferative neoplasms inferred from phylogenies. Nature, 602(7895):162–168, 2022.

9. Henry Lee-Six, Nina Friesgaard Øbro, Mairi S Shepherd, Sebastian Grossmann, Kevin Dawson, Miriam Belmonte, Robert J Osborne, Brian JP Huntly, Inigo Martincorena, Elizabeth Anderson, et al. Population dynamics of normal human blood inferred from somatic mutations. Nature, 561(7724):473–478, 2018.

10. Martin A Nowak, Franziska Michor, and Yoh Iwasa. The linear process of somatic evolution. Proceedings of the national academy of sciences, 100(25):14966–14969, 2003.

11. Timour Baslan and James Hicks. Unravelling biology and shifting paradigms in cancer with single-cell sequencing. Nature Reviews Cancer, 17(9):557–569, 2017.

12. Nicholas Navin, Jude Kendall, Jennifer Troge, Peter Andrews, Linda Rodgers, Jeanne McIndoo, Kerry Cook, Asya Stepansky, Dan Levy, Diane Esposito, et al. Tumour evolution inferred by single-cell sequencing. Nature, 472(7341):90–94, 2011.

13. Tanja Stadler, Roger Kouyos, Viktor von Wyl, Sabine Yerly, Jürg Böni, Philippe Bürgisser, Thomas Klimkait, Beda Joos, Philip Rieder, Dong Xie, et al. Estimating the basic reproductive number from viral sequence data. Molecular biology and evolution, 29(1):347–357, 2012.

14. Tanja Stadler, Denise Kühnert, Sebastian Bonhoeffer, and Alexei J Drummond. Birth–death skyline plot reveals temporal changes of epidemic spread in hiv and hepatitis c virus (hcv). Proceedings of the National Academy of Sciences, 110(1):228–233, 2013.

15. Denise Kühnert, Tanja Stadler, Timothy G Vaughan, and Alexei J Drummond. Simultaneous reconstruction of evolutionary history and epidemiological dynamics from viral sequences with the birth– death sir model. Journal of the Royal Society Interface, 11(94):20131106, 2014.

16. Alexandra Gavryushkina, David Welch, Tanja Stadler, and Alexei J Drummond. Bayesian inference of sampled ancestor trees for epidemiology and fossil calibration. PLoS computational biology, 10(12):e1003919, 2014.

17. John Frank Charles Kingman. The coalescent. Stochastic processes and their applications, 13(3):235– 248, 1982.

18. Robert C Griffiths and Simon Tavaré. Ancestral inference in population genetics. Statistical science, pages 307–319, 1994.

19. John Wakeley. Coalescent theory: an introduction. Roberts & Co. Publishers, Greenwood Village, Colo., 2009.

20. Robert C Griffiths. Lines of descent in the diffusion approximation of neutral wright-fisher models. Theoretical population biology, 17(1):37–50, 1980.

21. M Nordborg, DJ Balding, M Bishop, and C Cannings. Handbook of statistical genetics. John Wiley & Sons, Chichester, UK, pages 179–212, 2001.

22. Richard R Hudson. Gene genealogies and the coalescent process. Oxford surveys in evolutionary biology, 7(1):44, 1990.

23. Robert C Griffiths and Simon Tavare. Sampling theory for neutral alleles in a varying environment. Philosophical Transactions of the Royal Society of London. Series B: Biological Sciences, 344(1310):403–410, 1994.

24. Peter Donnelly and Simon Tavare. Coalescents and genealogical structure under neutrality. Annual review of genetics, 29(1):401–421, 1995.

25. Allen G Rodrigo and Joseph Felsenstein. Coalescent approaches to hiv population genetics. The evolution of HIV, pages 233–272, 1999.

26. Korbinian Strimmer and Oliver G Pybus. Exploring the demographic history of dna sequences using the generalized skyline plot. Molecular Biology and Evolution, 18(12):2298–2305, 2001.

27. Alexei J Drummond, Andrew Rambaut, BETH Shapiro, and Oliver G Pybus. Bayesian coalescent inference of past population dynamics from molecular sequences. Molecular biology and evolution, 22(5):1185–1192, 2005.

28. Erik M Volz, Sergei L Kosakovsky Pond, Melissa J Ward, Andrew J Leigh Brown, and Simon DW Frost. Phylodynamics of infectious disease epidemics. Genetics, 183(4):1421–1430, 2009.

29. Oliver G Pybus and Andrew Rambaut. Genie: estimating demographic history from molecular phylogenies. Bioinformatics, 18(10):1404–1405, 2002.

30. Alexei J Drummond, Geoff K Nicholls, Allen G Rodrigo, and Wiremu Solomon. Estimating mutation parameters, population history and genealogy simultaneously from temporally spaced sequence data. Genetics, 161(3):1307–1320, 2002.

31. Christian P Robert. A note on jeffreys-lindley paradox. Statistica Sinica, pages 601–608, 1993.

32. David Madigan and Adrian E Raftery. Model selection and accounting for model uncertainty in graphical models using occam’s window. Journal of the American Statistical Association, 89(428):1535– 1546, 1994.

33. Bradley P Carlin and Siddhartha Chib. Bayesian model choice via markov chain monte carlo methods. Journal Of The Royal Statistical Society Series B: Statistical Methodology, 57(3):473–484, 1995.

34. Robert E Kass and Adrian E Raftery. Bayes factors. Journal of the american statistical association, 90(430):773–795, 1995.

35. Edward C Holmes. The evolution and emergence of RNA viruses. Oxford University Press, 2009.

36. Julie L Lockwood, Martha F Hoopes, and Michael P Marchetti. Invasion ecology. John Wiley & Sons, 2013.

37. Mel Greaves and Carlo C Maley. Clonal evolution in cancer. Nature, 481(7381):306–313, 2012.

38. Ignacio A Rodriguez-Brenes, Natalia L Komarova, and Dominik Wodarz. Tumor growth dynamics: insights into evolutionary processes. Trends in ecology & evolution, 28(10):597–604, 2013.

39. Sébastien Benzekry, Clare Lamont, Afshin Beheshti, Amanda Tracz, John ML Ebos, Lynn Hlatky, and Philip Hahnfeldt. Classical mathematical models for description and prediction of experimental tumor growth. PLoS computational biology, 10(8):e1003800, 2014.

40. Xinyou Yin, JAN Goudriaan, Egbert A Lantinga, JAN Vos, and Huub J Spiertz. A flexible sigmoid function of determinate growth. Annals of botany, 91(3):361–371, 2003.

41. Benjamin Gompertz. Xxiv. on the nature of the function expressive of the law of human mortality, and on a new mode of determining the value of life contingencies. in a letter to francis baily, esq. frs &c. Philosophical transactions of the Royal Society of London, (115):513–583, 1825.

42. Elissa M Zullinger, Robert E Ricklefs, Kent H Redford, and Georgina M Mace. Fitting sigmoidal equations to mammalian growth curves. Journal of Mammalogy, 65(4):607–636, 1984.

43. Angela M Gibson, N Bratchell, and TA Roberts. The effect of sodium chloride and temperature on the rate and extent of growth of clostridium botulinum type a in pasteurized pork slurry. Journal of Applied Microbiology, 62(6):479–490, 1987.

44. Kathleen MC Tjørve and Even Tjørve. Shapes and functions of bird-growth models: how to characterise chick postnatal growth. Zoology, 113(6):326–333, 2010.

45. Robert E Ricklefs. Patterns of growth in birds. Ibis, 110(4):419–451, 1968.

46. Remco Bouckaert, Timothy G Vaughan, Jöelle Barido-Sottani, Sebastián Duchêne, Mathieu Fourment, Alexandra Gavryushkina, Joseph Heled, Graham Jones, Denise Kühnert, Nicola De Maio, Michael Matschiner, Fabio K Mendes, Nicola F Müller, Huw A Ogilvie, Louis du Plessis, Alex Popinga, Andrew Rambaut, David Rasmussen, Igor Siveroni, Marc A Suchard, Chieh-Hsi Wu, Dong Xie, Chi Zhang, Tanja Stadler, and Alexei J Drummond. Beast 2.5: An advanced software platform for bayesian evolutionary analysis. PLoS Comput Biol, 15(4):e1006650, Apr 2019.

47. Joseph Felsenstein. Evolutionary trees from dna sequences: a maximum likelihood approach. Journal of molecular evolution, 17:368–376, 1981.

48. Alexei J Drummond, Oliver G Pybus, Andrew Rambaut, Roald Forsberg, and Allen G Rodrigo. Measurably evolving populations. Trends in ecology & evolution, 18(9):481–488, 2003.

49. Anna K Laird. Dynamics of growth in tumors and in normal organisms. National Cancer Institute Monograph, 30:15–28, 1969.

50. Guy Baele, Philippe Lemey, Andrew Rambaut, and Marc A Suchard. Adaptive mcmc in bayesian phylogenetics: an application to analyzing partitioned data in beast. Bioinformatics, 33(12):1798– 1805, 2017.

51. Charles J Geyer. Markov chain monte carlo maximum likelihood. 1991.

52. Nicola F Müller and Remco R Bouckaert. Adaptive parallel tempering for beast 2. BioRxiv, page 603514, 2019.

53. Alexei J Drummond, Kylie Chen, Fabio K Mendes, and Dong Xie. Linguaphylo: A probabilistic model specification language for reproducible phylogenetic analyses. PLoS Comput Biol, 19(7):e1011226, Jul 2023.

54. Alexey Kozlov, Joao M Alves, Alexandros Stamatakis, and David Posada. Cellphy: accurate and fast probabilistic inference of single-cell phylogenies from scdna-seq data. Genome biology, 23(1):37, 2022.

55. Kylie Chen, Jiří C Moravec, Alex Gavryushkin, David Welch, and Alexei J Drummond. Accounting for errors in data improves divergence time estimates in single-cell cancer evolution. Molecular biology and evolution, 39(8):msac143, 2022.

56. Ray R Arthur, Nassef Farahat Hassan, Mahasan Yousef Abdallah, Mohamed Said El-Sharkawy, Magdy Darwish Saad, Barbara G Hackbart, and Imam Zaghloul Imam. Hepatitis c antibody prevalence in blood donors in different governorates in egypt. Transactions of the Royal Society of Tropical Medicine and Hygiene, 91(3):271–274, 1997.

57. Mostafa Habib, Mostafa K Mohamed, Fatma Abdel-Aziz, Laurence S Magder, Mohamed Abdel-Hamid, Foda Gamil, Salah Madkour, Nabiel N Mikhail, Wagida Anwar, G Thomas Strickland, et al. Hepatitis c virus infection in a community in the nile delta: risk factors for seropositivity. Hepatology, 33(1):248– 253, 2001.

58. Stuart C Ray, Ray R Arthur, Anthony Carella, Jens Bukh, and David L Thomas. Genetic epidemiology of hepatitis c virus throughout egypt. The Journal of infectious diseases, 182(3):698–707, 2000.

59. Christina Frank, Mostafa K Mohamed, G Thomas Strickland, Daniel Lavanchy, Ray R Arthur, Laurence S Magder, Taha El Khoby, Yehia Abdel-Wahab, Wagida Anwar, Ismail Sallam, et al. The role of parenteral antischistosomal therapy in the spread of hepatitis c virus in egypt. The Lancet, 355(9207):887–891, 2000.

60. OG Pybus, AJ Drummond, T Nakano, BH Robertson, and A Rambaut. The epidemiology and iatrogenic transmission of hepatitis c virus in egypt: a bayesian coalescent approach. Molecular biology and evolution, 20(3):381–387, 2003.

