## Supplemental file for "Bayesian model-averaging of parametric coalescent models for phylodynamic inference"

#### S.1 Gompertz-Laird Model Derivations

The Gompertz model, characterised by an exponentially decreasing growth rate, is widely used to describe tumour growth dynamics. Among the various parameterisations proposed in the literature, the Gompertz-Laird formulation [? ] is frequently adopted. This classical representation involves three parameters: the initial population size $N_0$, the growth rate parameter $b$, and the carrying capacity $N_\infty$. The model is given by:

$$N(t) = N_0 \exp\big(\log\big(\tfrac{N_\infty}{N_0}\big)(1 - \exp(-bt))\big).$$

While this parameterisation is conceptually straightforward, it may not be optimal for Bayesian inference. In general, parameter sets that are less correlated facilitate the specification of marginal priors and improve the efficiency of Markov Chain Monte Carlo (MCMC) sampling. The strong correlation between $N_0$ and $N_\infty$ can impede effective mixing, thus slowing convergence. To address this issue, adaptive methods such as the Adaptable Variance Multivariate Normal (AVMN) operator [? ] can be employed to learn and adjust parameter correlations during the MCMC run. Additionally, alternative parameterisations can be considered to further reduce correlation and enhance computational performance.

To enhance the flexibility of prior specification and reduce parameter correlations, we reparameterise the Gompertz model using $N_0$, $f_0$, and $b$, where $f_0 = \frac{N_0}{N_\infty}$. This ratio $f_0$ is dimensionless and constrained to the interval $[0, 1]$, enabling the straightforward use of a Beta prior without directly depending on the absolute sizes $N_0$ or $N_\infty$. Under this parameterisation, $N_0$ sets the initial population size, $b$ governs the time scale, and $f_0$ locates the population along its growth trajectory. The reparameterised Gompertz model is:

$$N(t) = N_0 \exp\big(\log\big(\tfrac{1}{f_0}\big)(1 - \exp(-bt))\big).$$

To facilitate a direct comparison with the logistic model, we further reparameterise the Gompertz model using the same set of parameters. Specifically, we replace $N_0$ with an expression involving $N_\infty$, the time $t_{50}$ at which the population reaches half of $N_\infty$, and the growth rate $b$:

$$N_0 = N_\infty \cdot 2^{-\exp(bt_{50})}.$$

This adjustment removes the explicit dependence on $N_0$, yielding:

$$N(t) = N_\infty \cdot 2^{-\exp(bt_{50})} \exp\big(\log\big(\tfrac{1}{2^{-\exp(bt_{50})}}\big)(1 - \exp(-bt))\big).$$

The Gompertz model can also be extended to incorporate an ancestral population size $N_A$. In this expanded version, the population grows from $N_A$ rather than approaching zero at large times. The Gompertz expansion function is:

$$N(t) = (N_0 - N_A) \exp\left(\log\left(\frac{N_\infty - N_A}{N_0 - N_A}\right)\big[1 - \exp(-bt)\big]\right) + N_A.$$

In the expanded model, the interpretation of $t_{50}$ is naturally adjusted. Specifically, $t_{50}$ now corresponds to the time at which the population reaches $N_A + \frac{N_\infty - N_A}{2}$, reflecting the shifted baseline introduced by the ancestral population size.

Under this definition, $N_0$ can be expressed as:

$$N_0 = (N_\infty - N_A)\exp\left(-\frac{\log(2)}{\exp(-bt_{50})}\right) + N_A.$$

### Supplementary Figures

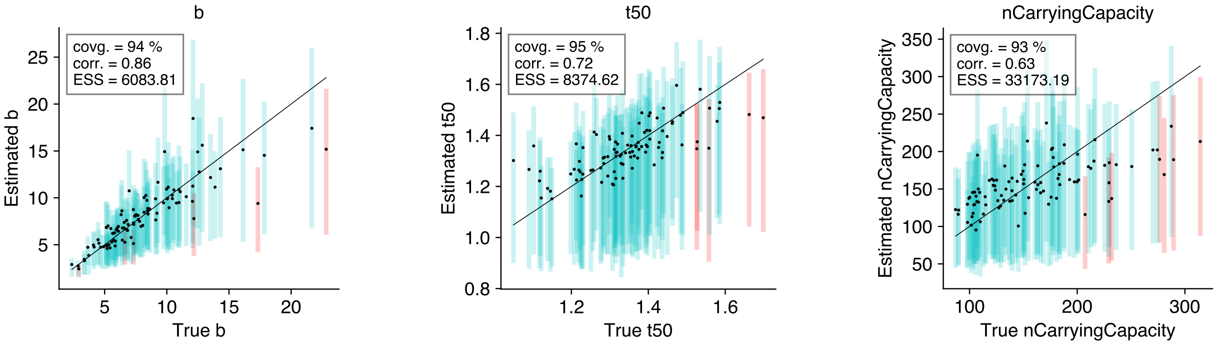


(a) Parameter 1 (b) Parameter 2 (c) Parameter 3

Supplementary Figure S.1: **Calibration results for the Logistic Coalescent model.** Each subfigure shows one of three key parameters inferred from 100 replicated simulations. The average ESS values for all parameters exceeded 200, indicating robust mixing and reliable inference.

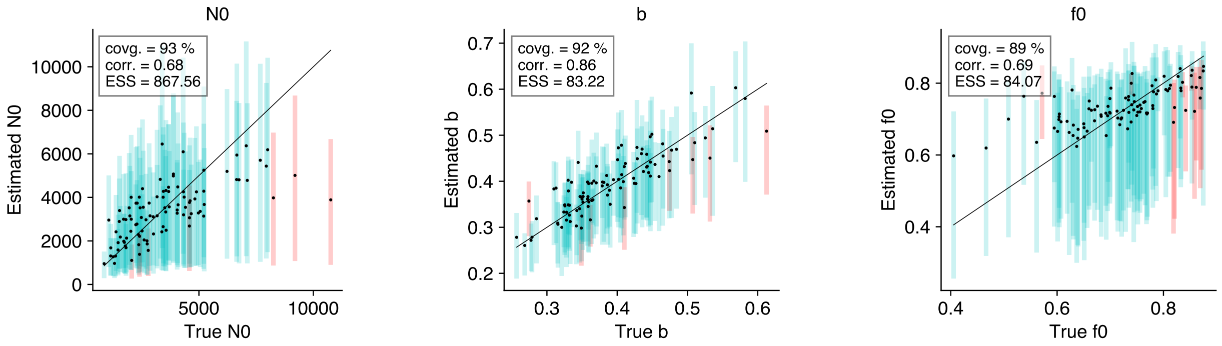


(a) N0 (b) b (c) f0

(d) First parameterization (top row)

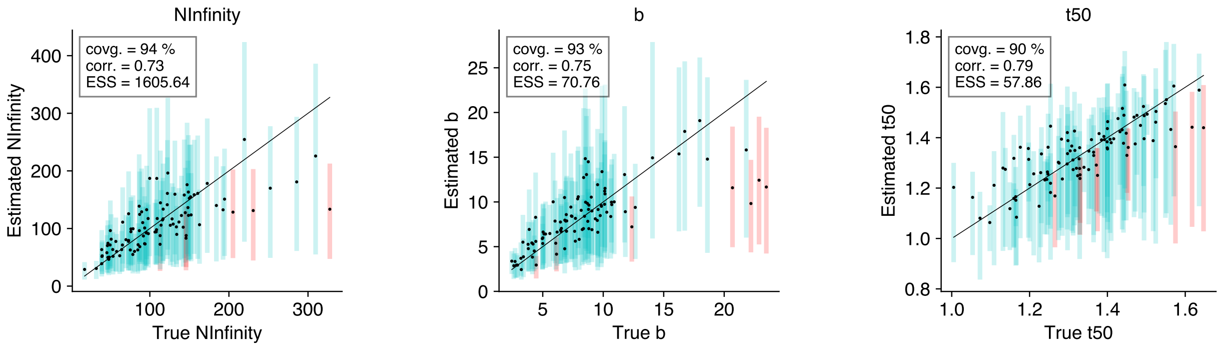


(e) NInfinity (f) b (g) t50

(h) Second parameterization (bottom row)

Supplementary Figure S.2: **Performance of the Gompertz Coalescent model under two parameterizations.** Using the same MCMC chain length as for the Logistic model resulted in substantially lower ESS values.

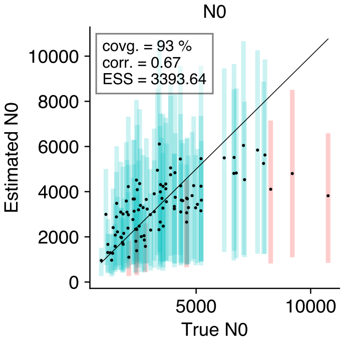


(a) N0

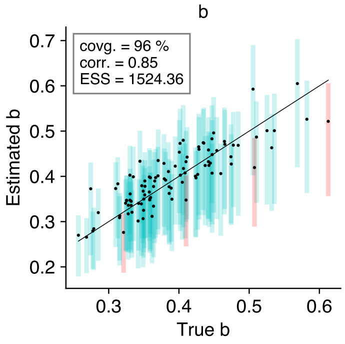


(b) b

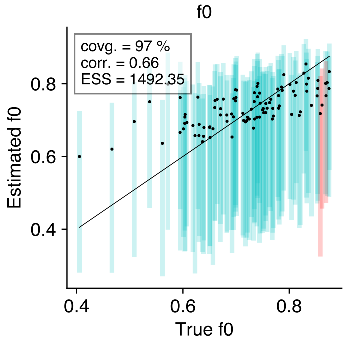


(c) f0

(d) First parameterization (top row)

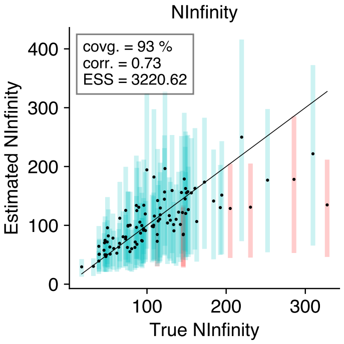


(e) NInfinity

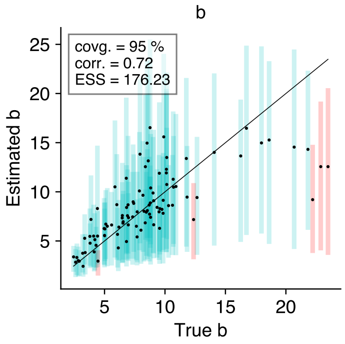


(f) b

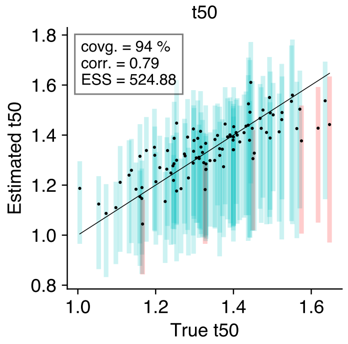


(g) t50

(h) Second parameterization (bottom row)

Supplementary Figure S.3: **Improvement in inference for the Gompertz Coalescent model after introducing specialized operators.** By incorporating up-down operators and the AVMN operator, the average ESS improved markedly. The figure shows 95% highest posterior density (HPD) intervals for key parameters, with blue indicating that the true value is captured and red indicating that it is not. After applying the new operators, 91%–99% of the true values fell within the estimated 95% HPD intervals, demonstrating significantly improved calibration.

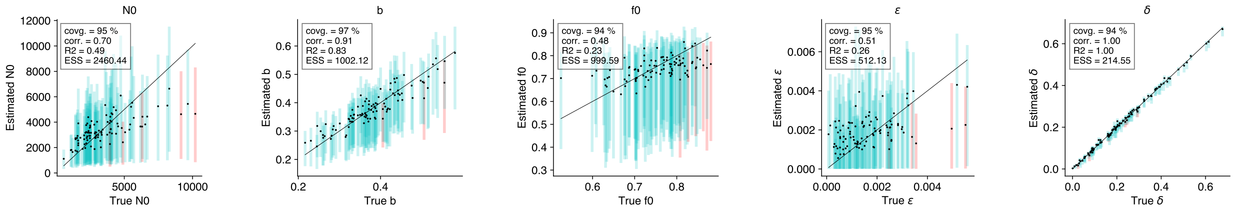


(a) N0 (b) b (c) f0 (d) $\varepsilon$ (e) $\delta$

(f) First parameterization (top row)

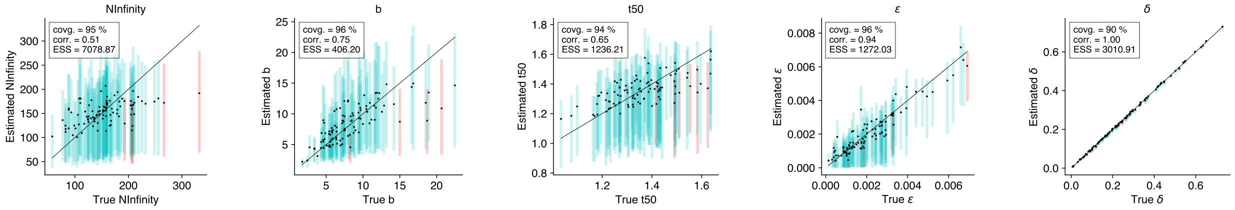


(g) NInfinity (h) b (i) t50 (j) $\varepsilon$ (k) $\delta$

(l) Second parameterization (bottom row)

Supplementary Figure S.4: **Calibration of the GT16 substitution and error model.** Even under increased complexity, the introduction of correlated-move operators and AVMN maintained satisfactory ESS values and reliable parameter estimates. The 95% HPD intervals contained 91%–99% of true parameter values, underscoring the robustness of our approach under complex scenarios.

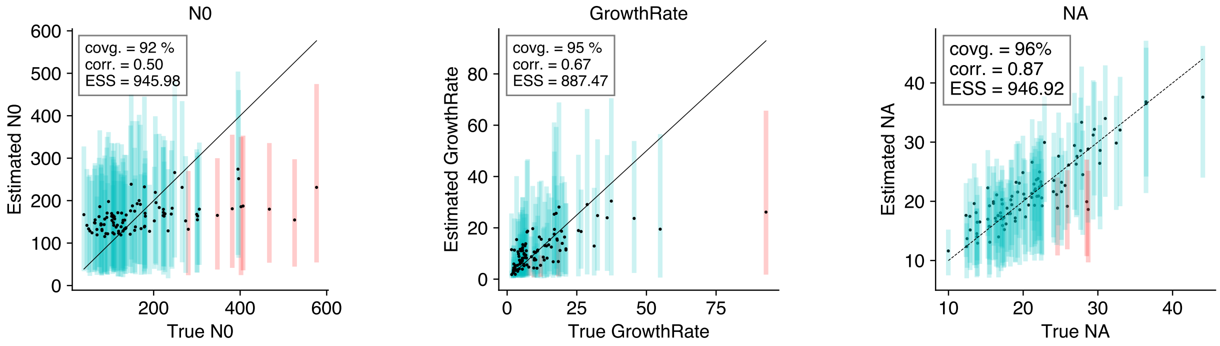


(a) $N_0$ (b) Growth rate $r$ (c) $N_A$

Supplementary Figure S.5: **Posterior estimation results for the three parameters of the exponential expansion model:** The initial population size $\mathbf{N_0}$, the growth rate $\mathbf{r}$, and the ancestral population size $\mathbf{N_A}$.

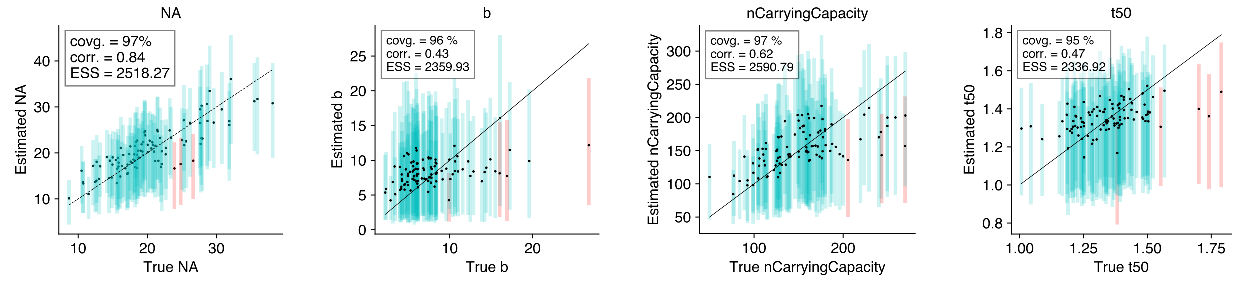


(a) $N_A$ (b) Growth rate $b$ (c) $n$CarryingCapacity (d) $t_{50}$

Supplementary Figure S.6: **Posterior estimation results for the four parameters of the logistic expansion model:** The four parameters of the logistic expansion model: (a) the ancestral population size $\mathbf{N_A}$, (b) the growth rate $\mathbf{b}$, (c) the carrying capacity **n**CarryingCapacity, and (d) the half-capacity time $\mathbf{t_{50}}$.

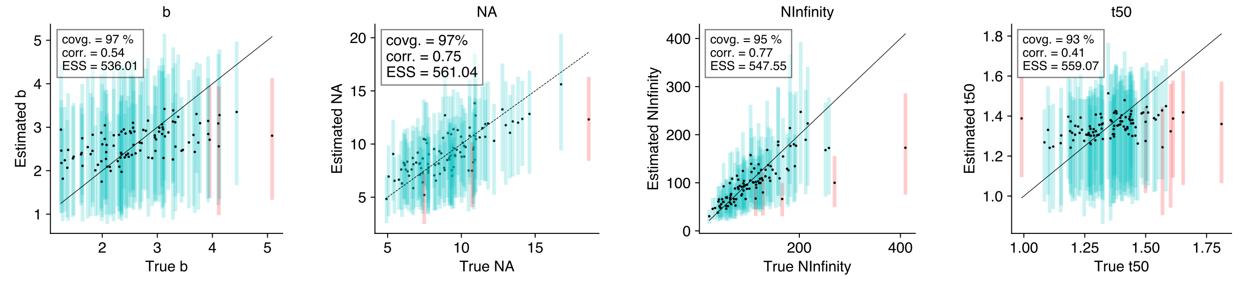


(a) Growth rate $b$ (b) $N_A$ (c) $N_\infty$ (d) $t_{50}$

Supplementary Figure S.7: **Posterior estimation results for the four parameters of the Gompertz expansion model (using the $t_{50}$ parameterization):** The growth rate $\mathbf{b}$, the ancestral population size $\mathbf{N_A}$, the carrying capacity $\mathbf{N_\infty}$, and the half time $\mathbf{t_{50}}$.

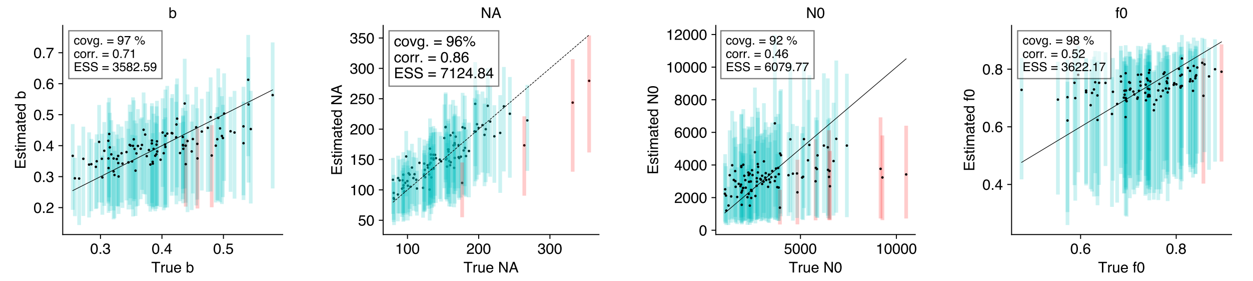


(a) Growth rate $b$ (b) $N_A$ (c) $N_0$ (d) $f_0$

Supplementary Figure S.8: **Posterior estimation results for the four parameters of the Gompertz expansion model (using the $\mathbf{f_0}$ parameterization):** The growth rate $\mathbf{b}$, the ancestral population size $\mathbf{N_A}$, the initial population size $\mathbf{N_0}$, and the dimensionless ratio $\mathbf{f_0}$.

#### S.1 LPhy Code for Model Averaging

```
model {
  π ~ Dirichlet(conc=[2.0, 2.0, 2.0, 2.0]);
  κ ~ LogNormal(meanlog=1.0, sdlog=0.5);
  b ~ LogNormal(meanlog=-2.2, sdlog=1.0);
  I_na ~ UniformDiscrete(lower=0, upper=1);
  N_max ~ LogNormal(meanlog=9.16, sdlog=2.0);
  NA ~ LogNormal(meanlog=5.0, sdlog=2.0);
  exponentialPopFunc = exponentialPopFunc(GrowthRate=b, N0=N_max, NA=NA, I_na=I_na);
  t50 ~ LogNormal(meanlog=3.9, sdlog=0.64);
  logisticPopFunc = logisticPopFunc(t50=t50, nCarryingCapacity=N_max, b=b, NA=NA, I_na=I_na);
  gompertzPopFunc = gompertzPopFunc_t50(t50=t50, b=b, NInfinity=N_max, NA=NA, I_na=I_na);
  x ~ Uniform(lower=10, upper=60);
  expansionPopFunc = ExpansionPopFunc(NA=NA, r=b, NC=N_max, x=x, I_na=I_na);
  tau ~ LogNormal(meanlog=4.6, sdlog=0.3);
  Cons_Exp_ConsPopFunc = Cons_Exp_ConsPopFunc(tau=tau, r=b, NC=N_max, x=x);

  }
```

Supplementary Code S.1: **LPhy script for analysing the HCV real dataset.** This script illustrates how multiple population dynamics models are combined through a stochastic variable selection approach to perform model averaging for the HCV dataset. Each model has its own parameters, drawn from specified prior distributions.

```
model {
  π ~ Dirichlet(conc=[2.0, 2.0, 2.0, 2.0]);
  κ ~ LogNormal(meanlog=1.0, sdlog=0.5);
  N_max ~ LogNormal(meanlog=4.5, sdlog=2.0);
  constantPopFunc = constantPopFunc(N0=N_max);
  b ~ LogNormal(meanlog=3, sdlog=1.0);
  I_na ~ UniformDiscrete(lower=0, upper=1);
  NA ~ LogNormal(meanlog=4.0, sdlog=2.0);
  exponentialPopFunc = exponentialPopFunc(GrowthRate=b, N0=N_max, NA=NA, I_na=I_na);
  t50 ~ LogNormal(meanlog=-2.2, sdlog=0.8);
  logisticPopFunc = logisticPopFunc(t50=t50, nCarryingCapacity=N_max, b=b, NA=NA, I_na=I_na);
  f0 ~ Beta(alpha=1, beta=1);
  gompertzPopFunc = gompertzPopFunc_f0(b=b, N0=N_max, f0=f0, NA=NA, I_na=I_na);

}
```

Supplementary Code S.2: **LPhy script for analysing the L86 real dataset.** This script illustrates how multiple population dynamics models are combined through a stochastic variable selection approach to perform model averaging for the L86 dataset. Each model has its own parameters, drawn from specified prior distributions.

```
model {
  π ~ Dirichlet(conc=[2.0, 2.0, 2.0, 2.0]);
  κ ~ LogNormal(meanlog=1.0, sdlog=0.5);
  N_max ~ LogNormal(meanlog=9.16, sdlog=2.0);
  constantPopFunc = constantPopFunc(N0=N_max);
  b ~ LogNormal(meanlog=-2.2, sdlog=1.0);
  I_na ~ UniformDiscrete(lower=0, upper=1);
  NA ~ LogNormal(meanlog=5.0, sdlog=2.0);
  exponentialPopFunc = exponentialPopFunc(GrowthRate=b, N0=N_max, NA=NA, I_na=I_na);
  t50 ~ LogNormal(meanlog=3.9, sdlog=0.64);
  logisticPopFunc = logisticPopFunc(t50=t50, nCarryingCapacity=N_max, b=b, NA=NA, I_na=I_na);
  gompertzPopFunc = gompertzPopFunc_t50(t50=t50, b=b, NInfinity=N_max, NA=NA, I_na=I_na);
  models = [constantPopFunc, exponentialPopFunc, logisticPopFunc, gompertzPopFunc];
  I ~ UniformDiscrete(lower=0, upper=length(models)-1);
  SVSmodel = stochasticVariableSelection(indicator=I, models=models);
  tree ~ CoalescentPopFunc(n=40, popFunc=SVSmodel);
  D ~ PhyloCTMC(L=500, Q=hky(kappa=κ, freq=π), mu=7.9E-4, tree=tree);
  height = tree.rootAge();
  length = tree.treeLength();
}
```

Supplementary Code S.3: **LPhy script for simulating a coalescent tree and sequence alignment under a mix of demographic models (Constant, Exponential, Logistic, and Gompertz).** In the code, the Constant model by definition does not include an expansion form, while the other three models optionally include a non-zero ancestral population (`NA`). The variable-selection indicator `I` enables switching among the four models in a single unified simulation.
